# Robust thermometry-imaging at sub-micrometer and millisecond-resolution by fluorescence lifetime microscopy allows for additional acquisition of multiple imaging channels

**DOI:** 10.64898/2026.06.18.733084

**Authors:** Bijeesh Meethale Mangalassery, Simon Fabiunke, Malte Schmick, Jan Huebinger

## Abstract

Temperature is a fundamental parameter governing all molecular processes, including those that define life. Fluorescence microscopy is a powerful tool to observe molecular processes in living systems in real time. Precise control and measurement of temperature during fluorescence microscopy is therefore essential. We present here a robust temperature measurement based on the excited-state lifetime of the widely available and relatively inexpensive fluorescent dye pentamethine cyanine (Cy5). The excited-state lifetime of Cy5 shows a monotonic decline in the measurement range of 0 °C – 80 °C. The measured dependency is linear until 39 °C and monoexponential above. The dependance of excited-state lifetime upon temperature is used to measure temperature up to a precision of 0.5 °C or less, a temporal resolution down to <1 millisecond and to resolve temperature gradients with spatial resolutions that are only diffraction-limited. The far-red excitation and emission of Cy5 leaves bandwidth to simultaneously measure at least 3 additional spectral channels in standard fluorescent microscopes simultaneously. We demonstrate determination of temperature during 4-color live-cell fluorescence microscopy for a temperature-controlled experiment. We also show its applicability in measuring temperature gradients and laser-induced sample heating such as during STED nanoscopy.

## Introduction

The advantages of temperature measurements by laser-induced fluorescence imaging have been described in a recent review article ^1^. It can be utilized in a broad range of applications including the measurement of flame temperatures ^2^, in flow fields, cryogenics, ice-water mixtures, combustion engines ^1^ and even inside living cells ^3,4^. In living samples, temperature is an important parameter representative of thermodynamics of the system and local temperature reflects metabolic turnover ^5^. As an example, fast growing tumors produce an increased amount of heat ^6^. Temperature is also an important parameter for fluorescence microscopy experiments, such as for cryobiology ^7^, cryo-fixation ^8–10^, temperature-controlled cellular endocytosis ^11^ or (bio-)chemical reactions ^12^ and local laser heating ^13^. Finally, it can be used to check for unwanted laser-heating, which can perturb an experiment especially under intense laser irradiation, such as in stimulated emission depletion nanoscopy or multi-photon excitation ^14,15^. Resistance thermometers or thermocouples are usually very bulky compared to the imaging volume. Therefore, they do not measure temperature precisely at the imaging volume and the spatial resolution is low. Additionally, they change the thermal and optical properties of the sample significantly. Infrared thermometry can be used to measure thermal radiation from surfaces, but not inside the sample. Therefore, there is a need for precise and spatially resolved temperature measurements within the sample that can be performed simultaneously with fluorescence microscopy.

In principle, temperature can be measured by fluorescence intensity using temperature-dependent changes in fluorescence quantum yield or absorption cross section. For example, Rhodamine B shows a temperature-dependent decline in quantum yield between 25°C and 90°C. In contrast, the quantum yield of fluorescein is reported to be largely unaffected by temperature and the change of fluorescein intensity is dominated by an increasing absorption cross section with temperature ^1^. However, intensity in fluorescence microscopy can be influenced by many factors, such as absorption, reflection, photobleaching or changes in laser power. These changes can be more impactful across different samples, rendering intensity-based measures most useful as relative measurements within the same sample. One solution for a more robust, quantitative temperature measurement is a ratiometric approach, where two different dyes are contrasted ^16,17^. If the dyes reacting equally to external factors, this approach becomes self-calibrating. However, such measurements reserve a wide range of the spectrum of light that is available for fluorescence microscopy. This minimizes the region of the spectrum available for parallel measurements. Frequently, samples contain additional absorptive molecules, such as hemoglobins, which further reduces the available part of the spectrum.

Temperature can also be derived from fluorescence lifetime using a single excitation wavelength, utilizing that quantum yield of several dyes is temperature-dependent ^3,18^. Temperature-dependence of this lifetime of the excited-state can be linked to change in non-radiative decay rates, such as collisional quenching, structural fluorophore changes (e.g. isomerization), internal conversion or intersystem crossing ^19,20^. The excited-state lifetime can be spatially resolved by fluorescence lifetime imaging microscopy (FLIM)^21,22^ and it is not subjected to the aforementioned changes in the sample and is therefore a robust temperature measurement that uses only a single excitation wavelength ^3,23–25^.

Rhodamine B is the most widely used dye for temperature measurements ^1^. However, its broad spectrum with excitation starting at ~450 nm and emission ranging to ~700 nm, leaves little room for additional fluorescence measurements ^1^ and is hampered in hemoglobin-containing samples by strong absorption below 600 nm ^26^. On the other hand, near-infrared thermometry based on rare-earth-based particles with excitation wavelength >800 nm ^23,25^ is not compatible with most fluorescence microscopes. Further, measuring a soluble dye rather than distinct particles provides spatially continuous measurements. Therefore, there is a need for temperature measurements by fluorescence lifetime using a far-red dye. However, the intensity of red-shifted rhodamine dyes like rhodamine 640 or sulforhodamine 640, shows little temperature dependence between 25 and 90°C ^1^. On the other hand, fluorescence quantum yields of carbocyanines dissolved in alcohols are inversely correlated with temperature. This includes hexamethylindodicarbocyanine ^27^, which is the fluorescent core group of the well-established far-red fluorescence dye pentamethine cyanine (Cy5). Cy5 in aqueous media also shows a temperature-sensitive trans-to-cis photoisomerization pathway and singlet-to-triplet transition ^28^. The latter can be expected to be also temperature sensitive. Both effects would reduce fluorescence quantum yield and therefore lifetime of the excited-state. The triple-sulfonated version of Cy5 – a water soluble and commercially available dye – is therefore an interesting candidate for lifetime-based temperature measurements in aqueous environments.

Here, we characterize the excited-state lifetime of sulfonated Cy5 in water and dilute aqueous media over a temperature range between 0 and 80 °C to assess its potential as a temperature sensor. With this as a calibration, we determine temperature directly within and across the fluorescent microscope’s field of view. We demonstrate spatio-temporal resolution and show simultaneous multi-color imaging.

## Results

Using a custom-built temperature control chamber ^8,29^ on a confocal laser scanning microscope (CLSM), the fluorescence lifetime of Cy5 in water showed a strictly monotonical decrease with increasing temperatures ranging from 0 to 80 °C. In the temperature range between 0 °C and 39 °C, the lifetime decreased linearly (r^2^>0.9998; Suppl. Figure 1, Figure 1), whereas above 39 °C the temperature followed a monoexponential decay (r^2^=0.9999; Suppl. Figure 2, Figure 1). This was determined with two different microscopes using several different analysis methods (Suppl. Figures 3+4+5, see Methods). The whole temperature regime can also be fitted with a double exponential, where the 2^nd^ exponent has a negative coefficient or with a cubic function (both r^2^=0.9999; Suppl. Figures 6+7) for ease of conversion between lifetime and temperature. The residuals of any of these fits are <5 × 10^−3^. From the uncertainty of the lifetime measurement in each experiment (2σ), the uncertainty of temperature measurement was determined to be ≤0.5 °C (Figure 1 e). Dissolving Cy5 in a cell culture buffer instead of pure water resulted in a similar overall relationship between excited-state lifetime and temperature, albeit with slightly different values (Suppl. Figure 8). This emphasizes the need to consider solvent properties for temperature measurements by fluorescence.

**Figure 1:**
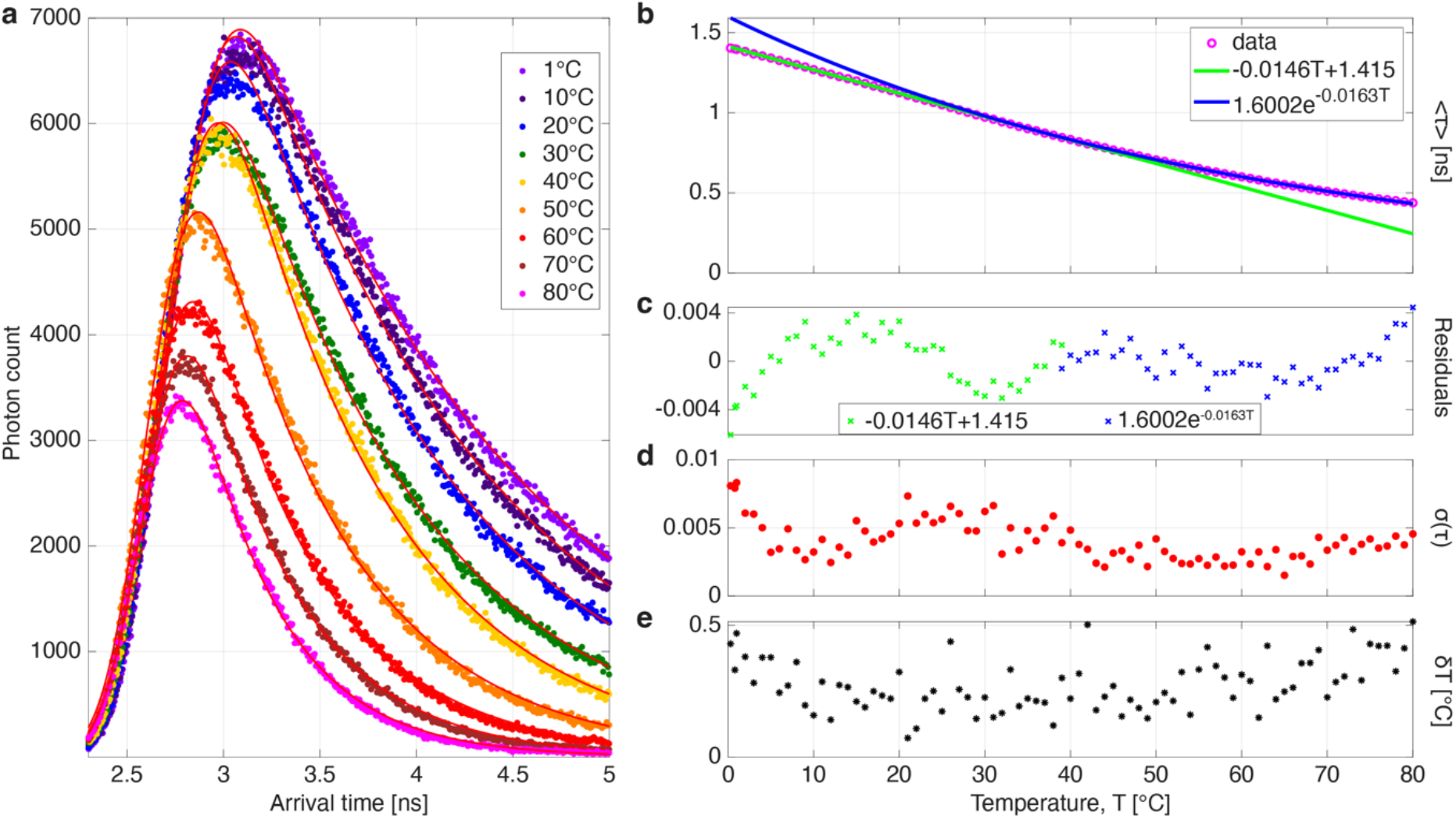
Excited-state lifetime of Cy5 as function of temperature. **a)** Exemplified lifetime histograms of Cy5 in water acquired on an Abberior Expert line CLSM at different temperatures (color code, upper right) individually fitted with monoexponential decay functions convolved with a gaussian approximation for the instrument response function. b) Mean excited-state lifetime (*τ*) of Cy5 as a function of temperature (N=4 independent experiments; cyan circles), fitted with a linear model over 0°C-39°C (green line) and a monoexponential model over 40°C-80°C (blue line). c) Residuals in both fit-regimes: linear (green symbols), monoexponential (blue symbols) d) Standard deviation (*σ*) of fitted lifetime across experiments (N=4). e) Uncertainty (*δ*) of temperature measurement calculated from 2*σ* in lifetime of individual experiments (N=4).

To induce temperature gradients in a sample on a fluorescence microscope, we utilized an infrared laser of 1470 nm to steadily illuminate a stationary spot of ∅ ≈ 24.4 μm full-width at half maximum (Figure 2a) in an aqueous sample containing Cy5 in a standard microscopy dish. A temperature-gradient that emanated from the illuminated spot by thermal dissipation was clearly measurable by the fluorescence lifetime distribution of Cy5 (Figure 2b, c). By collecting a sufficient number of photons per pixel, temperature gradients could therefore be resolved down to the diffraction-limit of the microscope.

**Figure 2:**
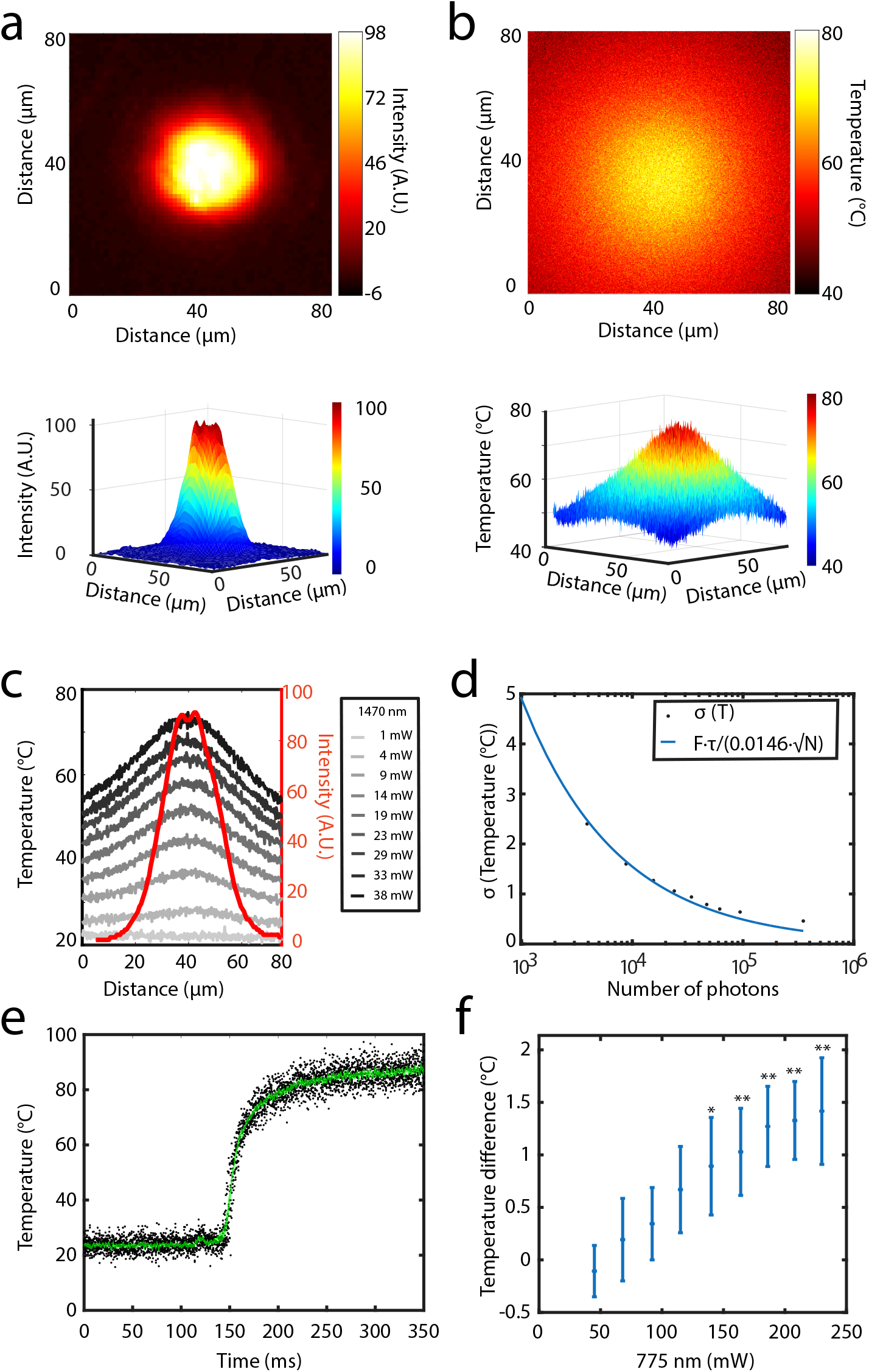
Sample heating by infrared lasers measured by Cy5 lifetime. a) Intensity profile of a stationary 1470-nm laser illumination of an aqueous Cy5 solution with 38 mW measured at the sample (≙ 8 × 10^3^ W/cm^2^) b) Corresponding temperature profile simultaneously measured by a scanning 640-nm laser. c) Temperature line-profiles measured at indicated laser powers of the 1470-nm lasers (black and grey lines) together with an exemplary intensity profile measured at 38 mW (red line). d) Standard deviation (*σ*, black dots) of the temperature measurement by fluorescence lifetime of Cy5 versus the number of photons per pixel (N). N was varied by binning pixels from an original image of 5 μm × 5 μm with 400 px x 400 px recorded at room temperature. Blue line: Data points fitted with the indicated function, combining the local slope of excited-state lifetime per 1 °C (0.0146, figure 1b, τ (25°C)= 1.05ns) and the dependence on shot noise 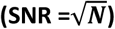 with the figure of merit of the fluorescence lifetime measurement (F=2.15). e) Temperature course upon turning-on 1470-nm laser illumination (110 mW), measured in the center of a flat-top laser beam of ∅ = 160 µm. Black dots: Accumulated data points from n=21 experiments; green lines: mean ± s.e.m. f) Temperature difference as a function of power of a highly focused, scanning 775-nm laser compared to minimal laser power (22 mW); N=10; *: p<0.05, **: p<0.01 compared to 22 mW

The accuracy of the measured lifetime, and thus of the derived temperature, scales inversely with the square root of the photon counts (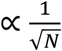, Figure 2d), as expected by the dependence on shot noise 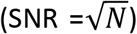^30^. The temperature can therefore be calculated with a standard deviation *σ*_*T*_ depending on the number of photons (N) and the figure of merit (F) with: 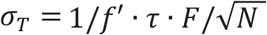, where *f*′ is the 1^st^ derivative of the function fitted to the measured excited-state lifetime over temperature (Figure 1b, Suppl. Figure 1-4, 6,7). The figure of merit 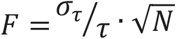^31^ was determined to be 2.15, by fitting measured *σ*_*T*_ over N (figure 2d). This also approximates a lower limit to the required number of photons per pixel for a diffraction-limited spatial measurement for a specific *σ*_*T*_. Consequentially, the temperature change induced by laser irradiation was measurable at a framerate of 200 µs with *σ*_*T*_ = 5.4 °C, or at a framerate of 1 ms with *σ*_*T*_ = 2.4 °C, at a single pixel level. By averaging 21 experiments, we could determine the temperature increase with a precision of ± 1.1 °C and ± 0.5 °C (s.e.m.), respectively. Even a short pre-pulse, which was also measurable using a photodiode 28 ms before the power increase, was detectable by a transient temperature increase (Figure 2e, Suppl. Figure 9).

The infrared irradiation wavelength of 1470 nm used to heat the sample is very close to a strong absorption peak of water with a linear absorption coefficient of ~3000 m^-1 32^. Using this wavelength, relatively low illumination densities of 8 × 10^3^ W/cm^2^ were enough to heat the sample to peak temperatures of >70 °C. We also measured the temperature inside the focus (~1.5 μm^2^) of a scanning, doughnut-shaped pulsed 775-nm laser, as frequently used for stimulated emission depletion (STED) nanoscopy. Since this pulsed STED laser perturbed the lifetime histogram of Cy5 when used in combination with 640-nm excitation, we fitted the histogram of fluorescence emanating from 775-nm excitation without an additional 640-nm pulse (Suppl. Figure 10). By this, a minor, yet measurable temperature increase was detectable (1.4 ± 0.5°C, mean ± s.d. at 230 mW ≙ 1.7 ×10^7^ W/cm^2^ vs 23 mW) within the moving spot excited by the laser beam in water (Figure 2f). This is consistent with a significantly lower absorption coefficient at this wavelength (~2.4 m^-1 33^) and with simulations for laser-induced sample heating of water samples under cryogenic conditions ^15^.

We then used Cy5 excited-state lifetime measurements to monitor temperatures in 4-color temperature-controlled fluorescence live-cell microscopy. For this, MCF7-cells were cultured on cover slides and attached to the temperature-controlled flow chamber. These cells expressed epidermal growth factor receptor fused to the fluorescent protein mCitrine (EGFR-mCitrine) and a phosphotyrosine binding domain fused to mCherry (PTB-mCherry) ^34,35^. They were incubated in cell culture medium containing Cy5 and cooled on-stage from 37°C to 5°C. Measuring the excited-state lifetime of Cy5 enabled determination of temperature in direct proximity to the cells (Figure 3a) and adjustment of the set temperature at the stage to the desired temperatures. In this experiment, an oil objective was used to obtain high resolution images of the cells. The oil made thermal contact between the microscope body at room temperature and the temperature-controlled flow chamber. In the custom-built temperature-controlled flow chamber we measured no significant temperature gradients across the field of view. However, an offset between the set and the measured temperature of ~1 to 3 °C was measured, likely due to heat exchange through the immersion oil. We then performed an EGF-binding experiment by incubating the cells in medium containing Dylight405-labeled EGF (360 ng/mL) at 5 °C. At this temperature, EGF binds to EGFR as apparent by the increase in fluorescence signal at the plasma membrane (Figure 3a, top row). However, receptor-mediated endocytosis is suppressed, which allows tracing of the endocytic process in a controlled way, upon heating back to 37 °C ^36^. After washing out excess EGF at 5 °C before elevating the temperature to 37 °C, endocytosis of EGF bound to EGFR could be clearly observed by the appearance of numerous fluorescence puncta co-occurring in the EGF and EGFR channels (Figure 3a). The EGF-induced phosphorylation of EGFR results in binding of PTB-mCherry as monitored by a decrease of mCitrine fluorescence lifetime upon Förster resonance energy transfer (FRET) between mCitrine and mCherry ^8,34,35^. From such data, the fraction of EGFR-mCitrine that exhibits FRET to PTBmCherry (*α*) can be computed in a global analysis by using *a priori* knowledge about the spatial invariance of the excited-state lifetime of the EGFR-mCitrine/PTB-mCherry complex and EGFR-mCitrine alone ^37–39^. Using this method, an increase in binding of PTB-mCherry to EGFR-mCitrine could be observed after return to 37°C, with subsequent dissociation of the EGFR-PTB-complex over time (Figure 3b). In endosomes however, PTB-binding remained high over the 60-min course of the experiment, as observable by their cyan color in figure 3b at 60 min. This is also corroborated by an increased PTB-mCherry signal at large endosomes taken at the end of the measurement (Figure 3a). It is of note that in this experiment, where the temperature was increased over 5 min from 5 °C to 37 °C, the early (5-min) phosphorylation response that is caused by (auto-)catalytic activation of EGFR at the plasma membrane ^40,41^ had a relatively low amplitude (Figure 3b).

**Figure 3:**
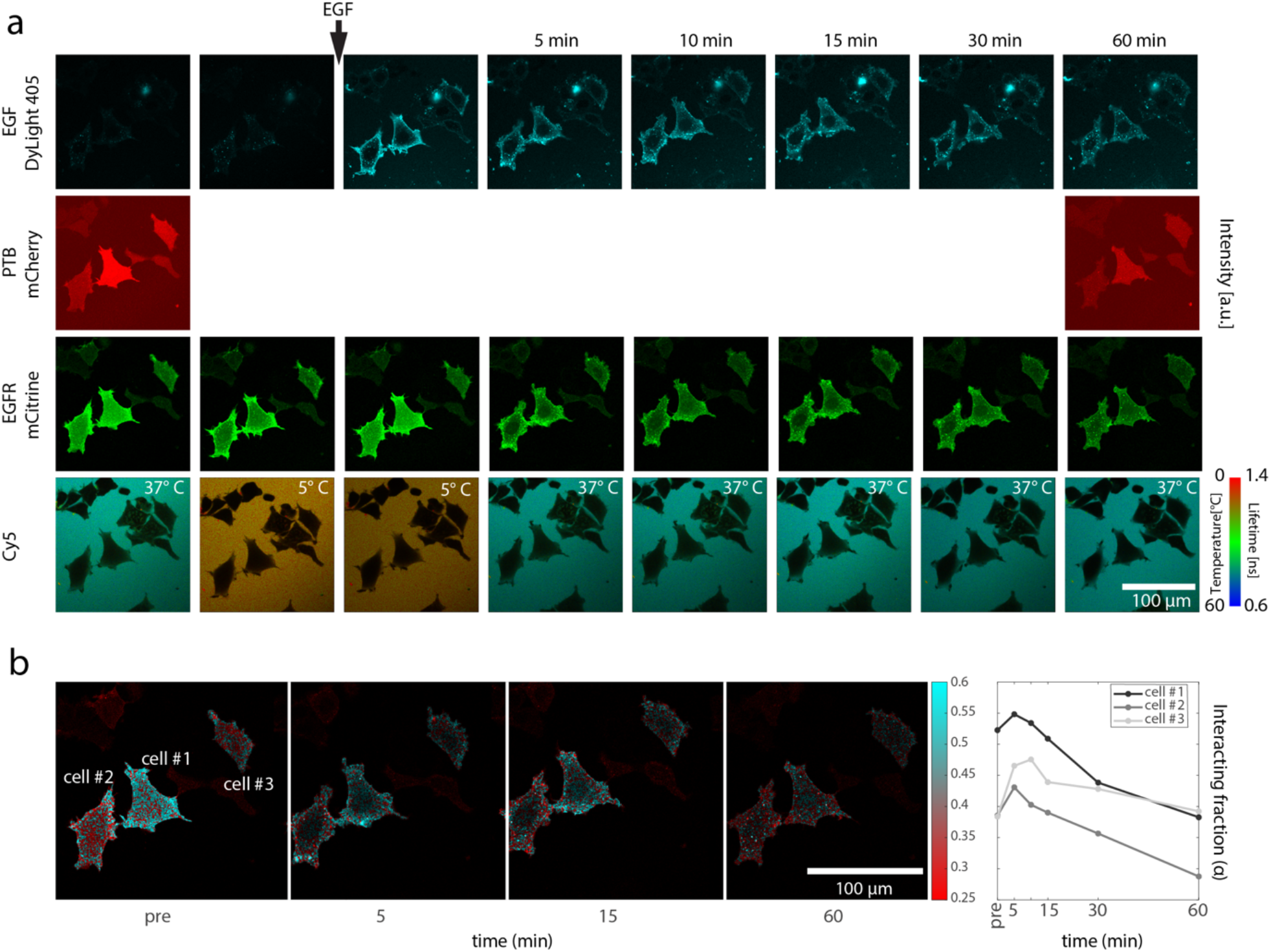
Temperature measurement in 4-color temperature-controlled live cell fluorescence microscopy. a) EGF-Dylight 405 (top row), PTB-mCherry (2^nd^ row) and EGFR-mCitrine (3^rd^ row) in MCF7 cells, together with temperature measurement by Cy5 excited-state lifetime (bottom row). Cells were cultured in a temperature-controlled flow-through chamber and first imaged at 37°C (first column), cooled to 5°C (2^nd^ column) incubated with 360 ng/mL EGF. After washing away excess EGF (3^rd^ column) the sample was heated back to 37°C and imaged at the indicated times after return to 37°C (4^th^ to 8^th^ column). b) Interacting fraction (α) of PTB-mCherry in complex with EGFR-mCitrine resulting in FRET measured by FLIM of mCitrine and determined in a global analysis. All photons of 3 masked cells across 6 micrographs at 37°C were accumulated to extrapolate a donor-only life-time (3.09 ns) and that of 100%-FRET (1.96 ns) by linear regression in a phasor plot. Left: spatial distribution of α at indicated times as normalized-intensity-weighted maps (colorbar). Right: cell-averaged α against time.

## Discussion

Using FLIM of Cy5 proved a robust and accessible way of determining temperature directly within microscopy samples with a temperature precision of ≤0.5 °C. This was tested over almost the entire range of liquid water (0 – 80 °C). Both overall temperatures and temperature differences in space and time could be extracted. Temporal resolution can be increased to below milliseconds for single pixel measurements on an APD-based TCSPC system. This could be further improved by using a fast lifetime contrast module ^42^, where more photons can be collected per time without disturbing the lifetime histogram. Spatial resolution was only limited by the diffraction limit of light in the microscopy setup.

We found a monotonical decrease of excited-state lifetime with temperature over the whole measurement range from 0 °C to 80 °C. Since the rate of fluorescence decay *k*_*f*_ is usually considered to be relatively constant with temperature, this decrease in excited-state lifetime most likely depends on competing rates such as the trans-to-cis isomerization rate (*k*_*ISO*_) and the singlet to triplet transition (*k*_*ISC*_) or other non-radiative decay rates (*k*_*nr*_) with:

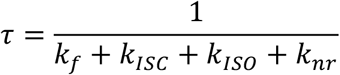

The trans-to-cis photoisomerization pathway of Cy5 in aqueous media has been shown to depend linearly on temperature in the temperature range between 11°C and 52°C ^28^. However, linear dependence on viscosity, as also shown in the same publication, would imply a flattening of the excited-state lifetime over temperature curve, due to the exponential decrease of viscosity on temperature. At room temperature *k*_*ISC*_ is of similar order as *k*_*ISO*_ for Cy5 ^28^. However, no data could be retrieved from the literature about the temperature dependence of *k*_*ISC*_. Also, for closely related carbocyanines dissolved in alcohols, a viscosity-dependent trans-to-cis photoisomerization was reported. Specifically, the fluorescence quantum yield of the Cy5-core molecule hexamethylindodicarbocyanine solved in ethanol also shows a clear trend to deviate from a linear relationship at higher temperatures ^27^. Here, we found a linear decrease in excited-state lifetime of Cy5 between 0 and 39°C followed by a monoexponetial decay between 40 and 80°C. This could in principle indicate a change in underlying non-radiative decay rates around the temperature of 40°C. However, it has to be taken into account that the shorter lifetimes at higher temperatures are closer to the width of the instrument response function (IRF), which increases the difficulty of distinguishing the excited-state lifetime from the IRF. Fitting of the IRF together with the fluorescence lifetime could in principle introduce a systematic bias. However, the same change was identified using two different microscopes with different IRFs and several different analysis methods. Importantly, knowledge about the exact photophysics is not relevant for the correlation between lifetime and temperature, as long as a reproducible lifetime-temperature calibration can be acquired. With such a calibration, FLIM of Cy5 can be used to measure real temperatures, including temperature gradients, directly across the microscopic field of view. Such a measurement can uncover a deviation to expected values or externally measured temperatures in a temperature-controlled fluidic device. Temperature can also be influenced by illumination, as demonstrated by temperature measurements during infra-red illumination, which showed drastic temperature increase of up to >70°C at relatively moderate illumination densities of 8 × 10^3^ W/cm^2^ across a spot of ∅ ≈ 24.4 μm. Absorption of a highly-focused, pulsed 775-nm laser beam, as used frequently for STED nanoscopy, did lead to a lesser, yet still measurable temperature of 1.4 ± 0.5°C increase in water despite much higher power density (1.7 ×10^7^ W/cm^2^ at 230 mW). This can be explained by a 3 orders of magnitude lower linear absorption coefficient of water at this wavelength. However, in material with a higher absorption coefficient, such as cells, a stronger temperature increase would be expected ^15^. In mouse brain, lasting tissue damage has been observed under focused illumination with 250 mW at 800 nm ^14^. The temperature increase will strongly depend on the sample geometry, e.g. thin samples plunge-frozen and imaged on electron microscopy grids for correlative cryo-fluorescence and electron microscopy heat dramatically more strongly, due to the inefficient heat dissipation in these samples ^15^.

The far-red excitation and emission of Cy5 allows simultaneous imaging of at least 3 other fluorescence colors. In 4-color fluorescence live-cell microscopy, temperature could be monitored precisely during an experiment, where receptor-mediated endocytosis was temporarily blocked by lowering temperature to 5°C and released again by heating back to the physiological temperature of 37°C. In the future, this will allow temperature block experiments at different precisely-defined temperatures directly on a fluorescence microscope. In this manner, endocytosed receptors could be directed to different endocytic compartments ^43^. It has been previously shown that signaling of EGFR from the plasma membrane and endocytic compartments can trigger different phenotypic outcomes induced by the same ligands ^41,44^ and that travelling through different compartments determines the signaling duration ^8,40^. Therefore, more detailed studies of interactions on and signaling from different compartments might reveal more of the intricacies of cellular signal transduction.

Temperature can be also measured by excited-state lifetime within living cells, which can reveal thermal inhomogeneities within the cells ^3,4^. However, measurements in cells are complicated by the inhomogeneous chemical and physical intracellular environment ^45^ and some results have been met with a degree of skepticism ^46,47^. Most recent investigations suggest a temperature dissipation slower than thermal conduction in living cells ^42^. Temperature gradients around the cells would not be captured by such measurements. Measurements in the homogeneous medium around the cells therefore provide a more robust measurement of temperature as an external parameter, whereas measurements in cells complementarily measure cellular properties.

Together, the presented results indicate a wide variety of potential applications in various fields, such as cell biology, (bio-)chemistry, cryobiological, material science and physics, where adding the widely available and relatively budget-friendly dye Cy5 to fluorescence microscopy experiments yields reproducible and spatially resolved temperature determination via FLIM.

## Methods

### Fluorescence Microscopy

Fluorescence microscopy was performed on I) an Abberior Expert line CLSM equipped with a 640-nm pulsed laser for excitation and a doughnut-shaped 1.25 W pulsed 775-nm depletion laser, an avalanche photon detector and a TCSPC FLIM system (Abberior Instruments GmbH, Göttingen, Germany) and an LCPLN100XIR (NA 0.85) objective, an UPlanSApo 100x/1.4NA oil immersion objective as well as an 40x/0.95NA LUCPLanXApo objective (Evident Europe GmbH, Hamburg, Germany) or II) on a Leica TSC SP8 CLSM (Leica Microsystems, Wetzlar, Germany), equipped with a fast lifetime contrast module (FALCON, Leica Microsystems), a 405-nm diode laser and a white light laser (white light laser Kit WLL2, NKT Photonics). Imaging was done with HC PL APO CS2 63x/1.4NA oil immersion objective. Following excitation wavelengths were used: DyLight 405 (405 nm), mCitrine (514 nm), mCherry (561 nm), Cy5 (640 nm). Fluorescence emission was detected by hybrid detectors (HyD) restricted at: DyLight 405 (415–460 nm), mCitrine (525–555 nm), mCherry (575–630 nm), Cy5 (655–720 nm).

### Analysis of Cy5 excited-state lifetimes from FLIM photon streams

Photon arrival times acquired on the TCSPC card of the Abberior expert line microscope were accumulated into lifetime histograms using a custom MATLAB script. Except for 775-nm excitation, the presented data was fitted using a gaussian approximation for the instrument response function that was obtained from the higher harmonic multiples of the laser frequency ^39^ convoluted with a monoexponential decay function. For spatial determination of temperature, photons out of the photon streams were mapped back into an image using their global arrival times. The parameters for the fit of the IRF were obtained globally from all accumulated photons to then only fit the fluorescence decay τ of the arrival time histograms on a per-pixel basis. A 5-parameter Gaussian with asymmetric flanks as an approximation for the IRF ^48^ showed the same linear and monoexponential regimes in the lifetime over temperature plots with very similar slopes and a ~0.1 ns offset. This multi-parameter analysis was however more prone to erroneous fit parameters and was therefore not routinely used in this work to directly fit fluorescence decay (Suppl. Figure 5). However, for the broader and strongly asymmetric pulse of the 775-nm laser the monoexponential decay was not fitted properly using the gaussian approximation for the IRF. Here, the more sophisticated model ^48^ was used to fit the IRF, which was measured by reflection from a glass slide. The obtained parameters were fixed for the convolution with a monoexponential decay that was then fitted to individual lifetime histograms.

Lifetime histograms acquired on the Leica Sp8 microscope were analyzed using the inbuilt software and a monoexponential decay function.

### Measurement of fluorescence lifetime over temperature

Temperature control was achieved with a custom-built temperature control stage that was described elsewhere in detail ^8,29^. Briefly, it consists of an aluminum block where a PID controller can adjust temperature by regulating flow of liquid nitrogen versus electric heating. A standard microscopy cover slide is attached to the block. Recesses in the aluminum block allow for media flow-through. Here the stage was used without its flow-through option and approximately 100 μL of 6 μM Cy5 in water were manually pipetted into the stage. An external 50-μm thermocouple was inserted into the temperature-controlled chamber to measure temperature directly within the sample. The thermocouple junction was imaged to precisely locate it, and fluorescence was collected from the immediately adjacent region to measure accurate local temperature. The openings of the chamber were closed by an adhesive (Blu Tack, Bostik, Puteaux, France) to avoid evaporation. At the Abberior Expert line CLSM, the sample was scanned with 400 × 400 (640-nm excitation) or 200 × 200 (775-nm excitation) pixels of 200 nm pixel length with a pixel dwell time of 200 μs. Care was taken to collect photons at a frequency <10 % of the laser repetition rate (40 MHz). The photon stream was collected until ~100 photons per pixel were collected (on average 5 frames). At the Leica SP8 CLSM, the sample was scanned with 512 × 512 pixel which covers an area of 184 μm × 184 μm, with a pixel size of 360 nm × 360 nm. A pixel dwell time of 102 μs was used and multiple frames are recorded until 200 photons per pixel were recorded. Lifetime was recorded under ascending and descending temperature, with no significant differences in the correlation between excited-state lifetime and temperature.

Uncertainties of the temperature measurement were calculated on individual experiments, which do not suffer from interexperimental uncertainties such as the uncertainty of the temperature measurement by the bare-wired thermocouple. For this, fluorescence lifetime was calculated on a per-frame basis of the respective accumulated photons. From this, mean and standard deviation (*σ*) of the fluorescence lifetime for each temperature was recorded. The uncertainty of the temperature measurement was calculated as:

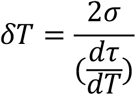

With *σ* being the standard deviation of the fluorescence lifetime (*τ*) measurement between frames.

### Local heating by infrared illumination

A continuous wave 1470-nm diode laser with maximum power of 30 W (Aerodiode, Bègles, France) was coupled in via the back port of the microscope and coupled into the beam path via a dichroic mirror. For spatial measurements (Figure 2 a-c), an aperture in the conjugated image plane was used to confine the illuminated area to 24.4 μm radius at full width and half maximum (Figure 2a). For temporal measurements (Figure 2 e), this aperture was omitted and the beam was scanned along 2 µm lines with a pixel size of 1 nm and 100 µs pixel dwell time in the center of the illuminated area. In this scanning region no temperature gradient was measurable. The time series was extracted by binning the photons according to their global arrival time. The resulting power of different voltages was measured after the microscope objective by a thermopile power sensor (Coherent Crop., Saxonburg, Pennsylvania, USA). A 100 μL sample of 6 μM Cy5 in water was imaged in an 8-well glass-bottom cell culture dish (Sarstedt, Nümbrecht, Germany) during illumination of 1470-nm laser light.

### 775-nm laser scanning illumination

The 775-nm illumination was realized on the Abberior Expert line microscope using laser scanning by the doughnut-shaped 775-nm laser. A 100 μL sample of 6 μM Cy5 in water was imaged in an 8-well glass-bottom cell culture dish (Sarstedt, Nümbrecht, Germany). The power of the 775-nm laser was measured after the objective by a thermopile power sensor (Coherent Crop., Saxonburg, Pennsylvania, USA). A calibration curve of lifetime over temperature was measured using the minimal power (24 mW) of the 775-nm laser that gave sufficient signal to obtain a lifetime histogram of sufficient signal over background (Suppl. Figure 10).

### Cell culture and transfection

MCF7 cells (86012803, ECACC) were cultured in Dulbecco’s modified eagle’s medium (DMEM; PAN-Biotech GmbH, Aidenbach, Germany) supplemented with 10% fetal bovine serum (FBS; PANBiotech GmbH), 2 mM L-Glutamine and 1% nonessential amino acids (NEAA, PAN-Biotech GmbH). The EGFR-mCitrine and PTB-mCherry plasmids were generated and validated previously ^35,40,41^.

One day before the experiment, cells were transfected by electroporation. For this, 10^6^ cells were washed twice in Opti-MEM medium (Thermo Fisher Scientific) and resuspended in 100 µL Opti-MEM medium containing 5 µg of each plasmid. The cell suspension was electroporated in an NEPA21 electroporator (Nepa Gene Co., Ltd., Ichikawa, Japan) using the standard settings of the manufacturer for MCF7 cells (2 poring pulses 125 V, 5 ms, 20 Hz; 5 transfer pulses 20 V, 50 ms, 20 Hz). Cells were resuspended in 1 mL DMEM + 10% FBS + 2 mM L-Glutamine + 1% NEAA + 100 μg/mL streptomycin, 100 U/mL penicillin.

### 4-color live-cell fluorescence microscopy with temperature control

5 × 10^5^ cells were seeded per 21 × 26 mm cover slide (Gerhard Menzel GmbH, Braunschweig, Germany) equipped with 10-μm double-sided sticky tape (Modulor GmbH, Berlin, Germany) in 6-well plates, as described before ^8,29^. In the center of the double-sided tape, a 21 × 5 mm rectangular area had been cut out before. Before the experiment, the cover slide was taped to the aluminum block of the temperature-controlled flow-chamber, which was assembled at the Leica SP8 microscope. The temperature was initially set to 37 °C and the chamber containing the cells was perfused with cell culture medium containing 6 μM Cy5 by an automated syringe system, which was connected by flexible tubes to the flow through chamber ^8,29^. The measurement in close proximity to the cells using the excited-state lifetime of Cy5 was used to adjust temperature to 37 °C at the cells. The sample was then cooled down to 5 °C, as confirmed by Cy5-lifetime next to the cells. Afterwards, the setup was perfused with cell culture medium containing 360 ng/mL EGF-DyLight 405. After 10 min, the cells were washed with fresh cell culture medium without EGF. After an additional 5 min of incubation, temperature was increased back to 37 °C within a time-course of 5 min.

### Quantification of FRET between mCitrine and mCherry from FLIM photon streams

mCitrine was excited using the white light laser at a frequency of 40 MHz and wavelength of 514 nm, and fluorescence emission was collected between 525 to 555 nm on a HyD. The donor (mCitrine) photon count images were thresholded to exclude background fluorescence. Assuming a fractional mixing of two species – donor-only (DO) EGFR-mCitrine vs. donor-acceptor complex (DA) allowing FRET to PTB-mCherry to occur – global analysis was performed as described previously by a custom MATLAB script to infer the spatial distribution of this interacting fraction α ^37–39,49^. For this, the accumulated distribution of photon-arrival-times of a diamond-shaped neighborhood of 12 pixel (diameter: 5 pixel) was converted into a phasor-plot of frequency-domain phase- and modulation-lifetime by Fourier transformation. Linear regression of the resulting distribution was used to globally extrapolate a DO lifetime (3.09 ns) and a DA lifetime (1.96 ns). α was calculated by projecting each 12N-pixel-neighborhood onto the line connecting DO and DA lifetime in the phasor plot and converted into a spatial map. This map was weighted by the normalized intensity map of per-pixel number-of-photons.

### Statistical analysis

Statistical testing was performed using GraphPad Prism 11.0.1 software. Columns were compared using repeated measures one-way ANOVA with Geisser-Greenhouse correction and Dunnett’s multiple comparison test.

## Data availability statement

All data generated or analyzed during this study are included in this published article (and its Supplementary Information files).

## Code availability

The MATLAB code used in this work is freely available under https://github.com/MPI-Dortmund/FLIM_Thermometry.

## Acknowledgments

The authors would like to thank Sabrina Seidler for excellent technical assistance in cell culture and transfection, Kirsten Michel for EGF-DyLight 405 and Roger Goody (all Max-Planck Institute of Molecular Physiology, Dortmund, Germany) for critical reading of the manuscript.

## Author contribution

JH conceived the project. BMM, SF and JH performed experiments. MS, BMM, JH and SF analyzed the data. JH wrote the manuscript with input from all authors.

