## Supplementary figures for "Robust thermometry-imaging at sub-micrometer and millisecond-resolution by fluorescence lifetime microscopy allows for additional acquisition of multiple imaging channels"

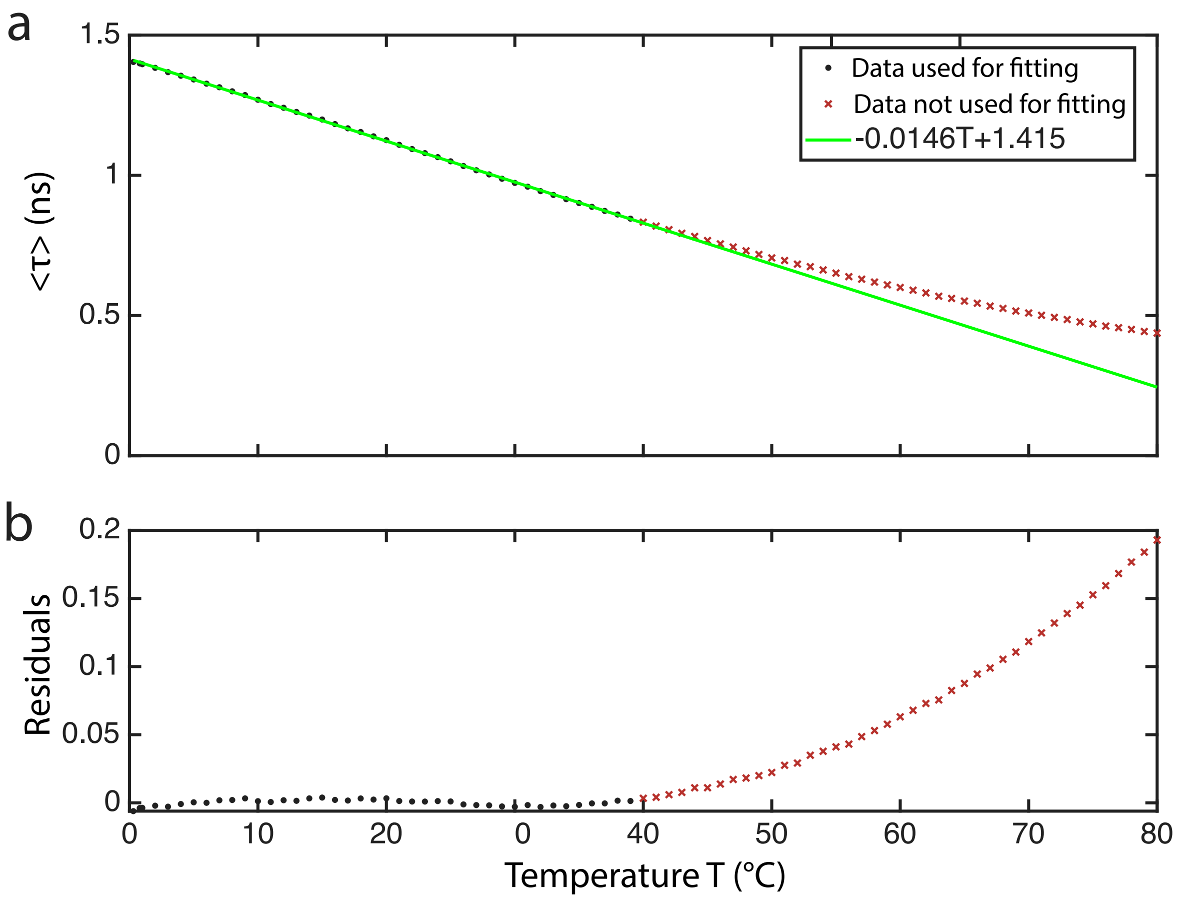


**Supplementary Figure 1: Linear regression to fluorescence lifetime of Cy5 vs temperature**

a) Plot of mean excited state lifetime ($\tau$) of Cy5 in water (black dots and red crosses) measured at an Abberior Expert line confocal laser scanning microscope against temperature (N=4) together with a linear function (green line) fitted to the data range from 0-39°C (black dots) b) Residuals from the temperature region included (black dots) and excluded (red crosses) for linear fitting.

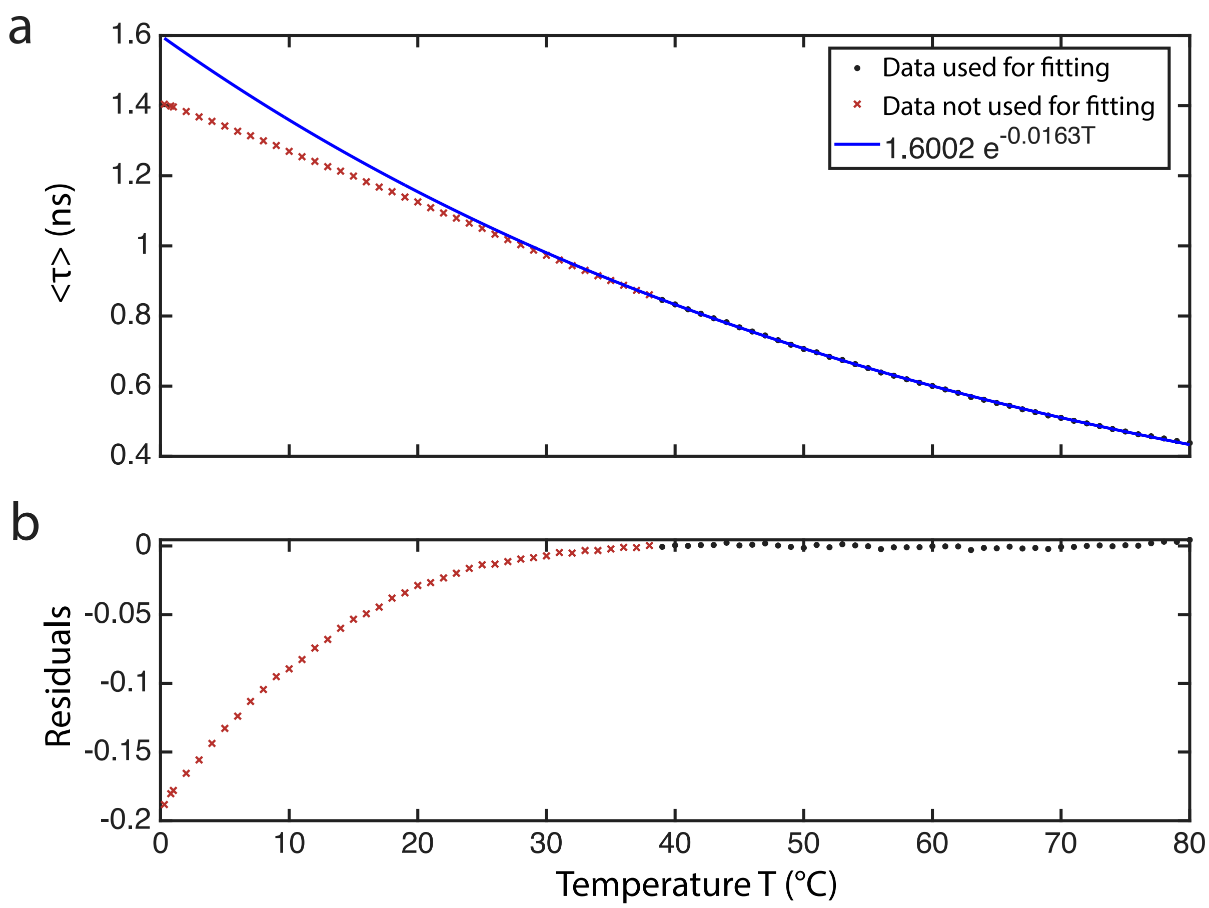


**Supplementary Figure 2: Monoexponential regression to fluorescence lifetime of Cy5 vs temperature**
a) Plot of mean excited state lifetime ($\tau$) of Cy5 in water (red crosses and black dots) measured at an Abberior Expert line confocal laser scanning microscope against temperature (N=4) together with a monoexponential decay function fitted (blue line) to the data range from 40-80°C (black dots) b) Residuals from the temperature region included (black symbols) and excluded (red symbols) for fitting with a monoexponential function.

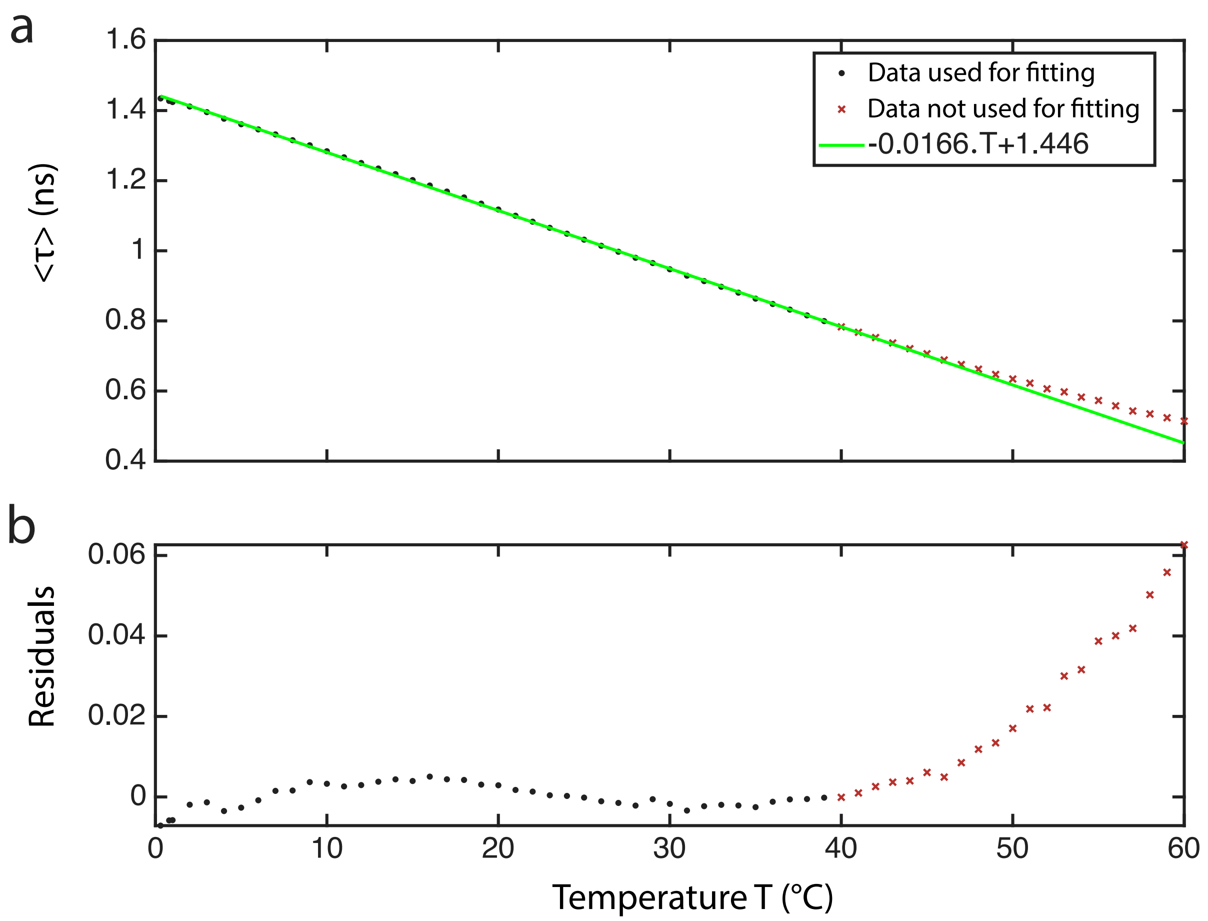


**Supplementary Figure 3: Linear regression to fluorescence lifetime of Cy5 vs temperature (different microscope)**

a) Plot of mean excited state lifetime ($\tau$) of Cy5 in cell culture medium (black dots and red crosses) measured at a Leica SP8 confocal laser scanning microscope against temperature (N=4) together with a linear function fitted (green line) to the data range from 0-39°C (black dots) b) Residuals from the temperature region included (black dots) and excluded (red crosses) for linear fitting.

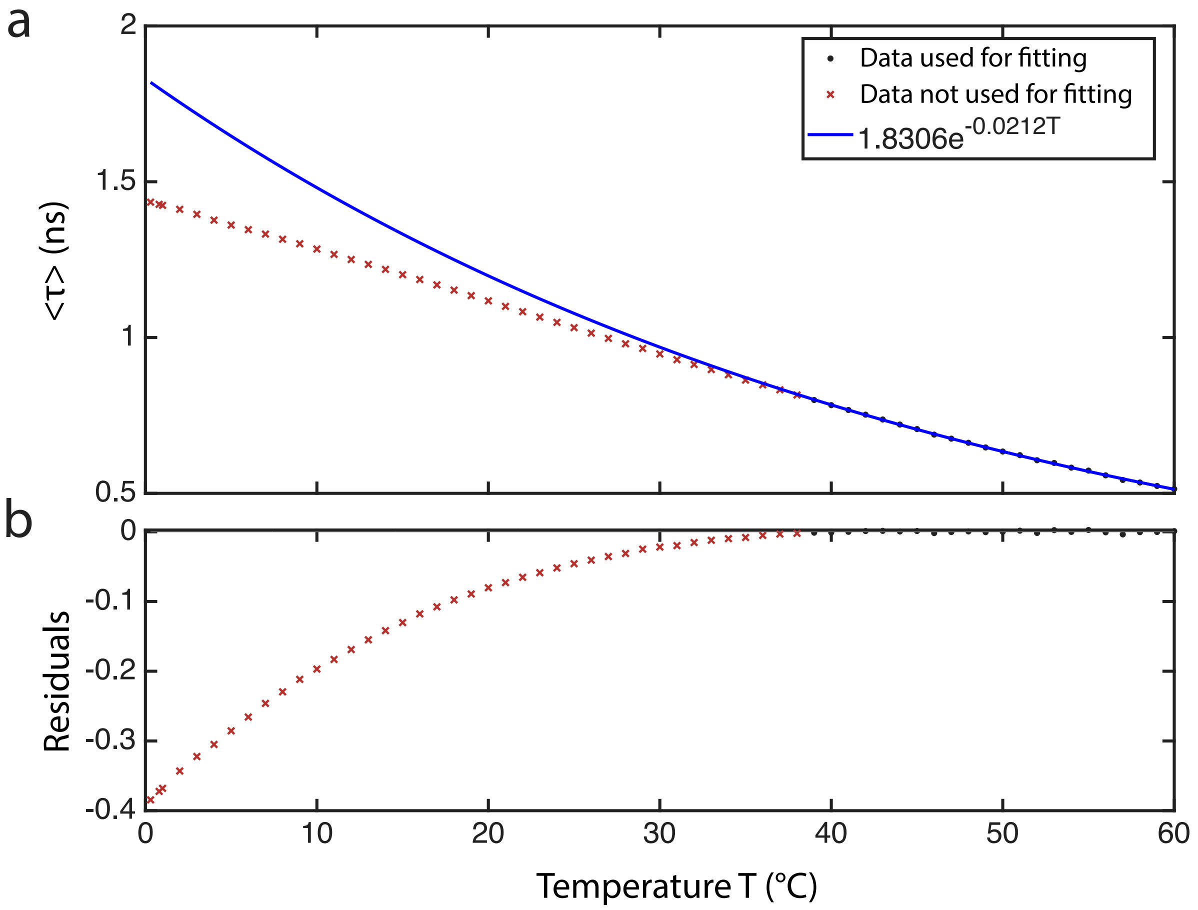


**Supplementary Figure 4: Monoexponential regression to fluorescence lifetime of Cy5 vs temperature (different microscope)**

a) Plot of mean excited state lifetime ($\tau$) of Cy5 in cell culture media (black dots and red crosses) measured at a Leica SP8 confocal laser scanning microscope against temperature (N=4) together with a monoexponential decay function fitted (blue line) to the data range from 40-60°C (black dots) b) Residuals from the temperature region included (black dots) and excluded (red crosses) for fitting with a monoexponential function.

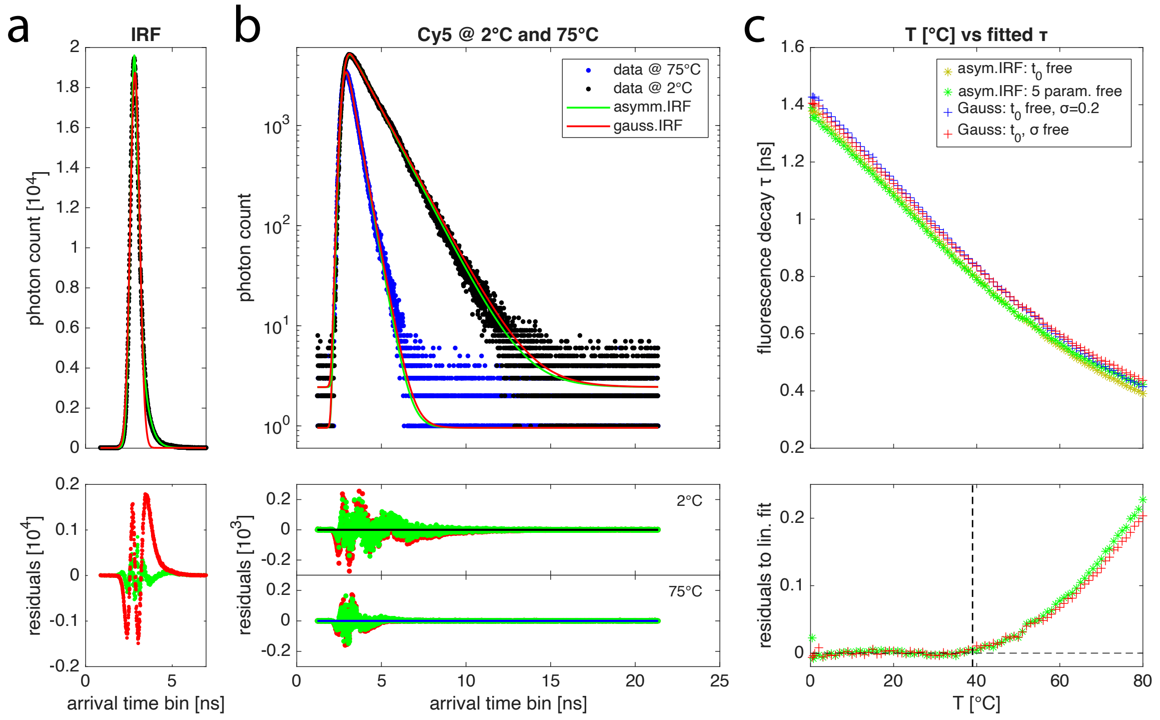


**Supplementary Figure 5: Impact of different IRF-models with parameters inferred from arrival-time histogram** a)-c): Top: measurement with fits, Bottom: respective residuals between data and respective fit. a) IRF measured as the histogram of number of photons of reflected pulsed 640-nm laser excitation (black dots) together with Gaussian fit (red line) and 5-parameter gaussian with asymmetric flanks (green line). b) Fit of a monoexponential fluorescence decay of Cy5-photon-arrival-times at 75°C (blue dots) and 2°C (black dots) individually convoluted with the two IRF-models in (a). c) Plot of fitted arrival times τ against externally controlled temperature T. Red: Fit with gaussian IRF described by parameters $t_0$ (center of Gaussian) and σ (width of Gaussian), free for each T; blue: using fixed σ=0.2 ns and free $t_0$ to fit the data; green: Fit with asymmetric IRF-model and individual 5 IRF-related parameters ($t_0$, and 4 parameters defining σ and asymmetric flanks) for each T; yellow: same as green, but fixing the 4 parameters defining σ and asymmetric flanks to those obtained at 39°C.

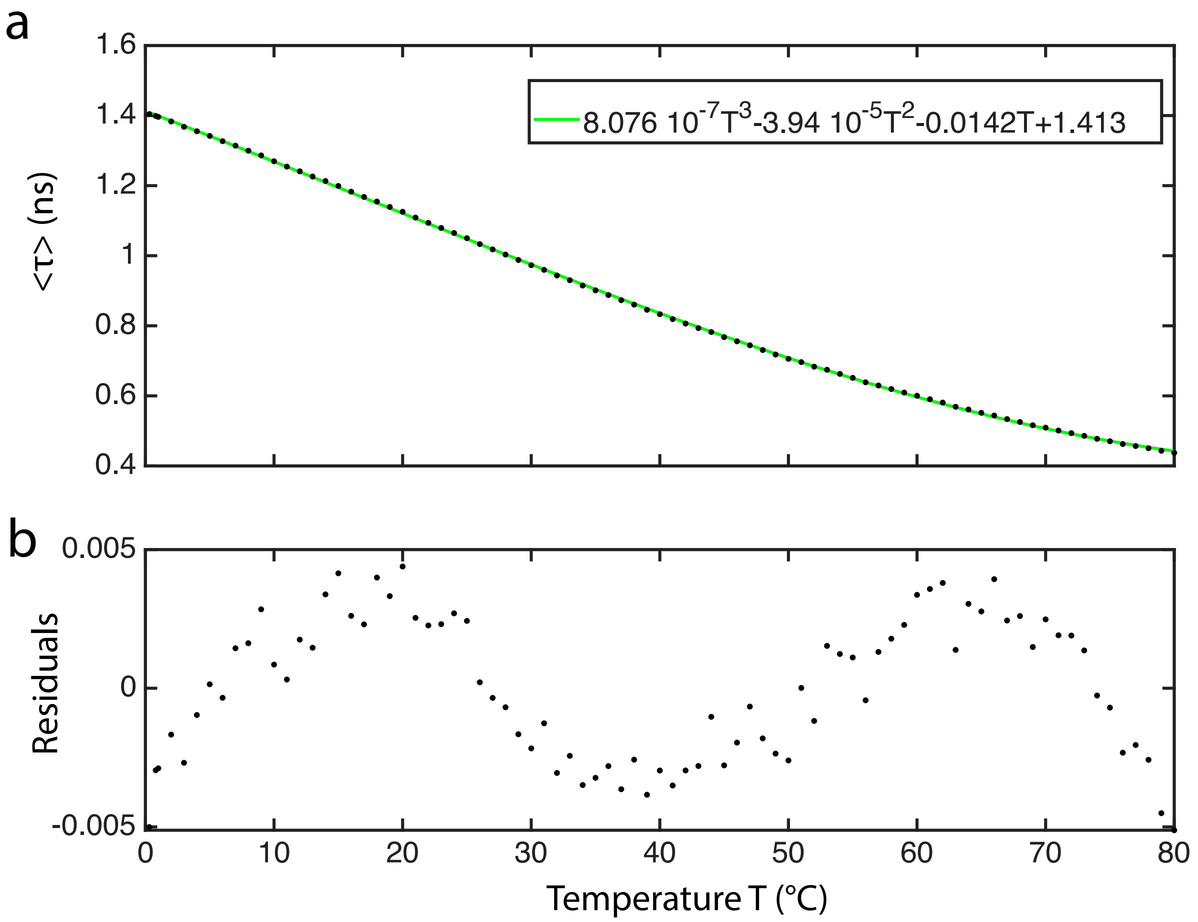


**Supplementary Figure 6: Cubic regression to fluorescence lifetime of Cy5 vs temperature**
a) Plot of mean excited state lifetime ($\tau$) of Cy5 in cell culture media (black dots) measured at an Abberior Expert line confocal laser scanning microscope against temperature (N=4) together with a cubic fit of the data (green line) b) Residuals from the fit (black dots).

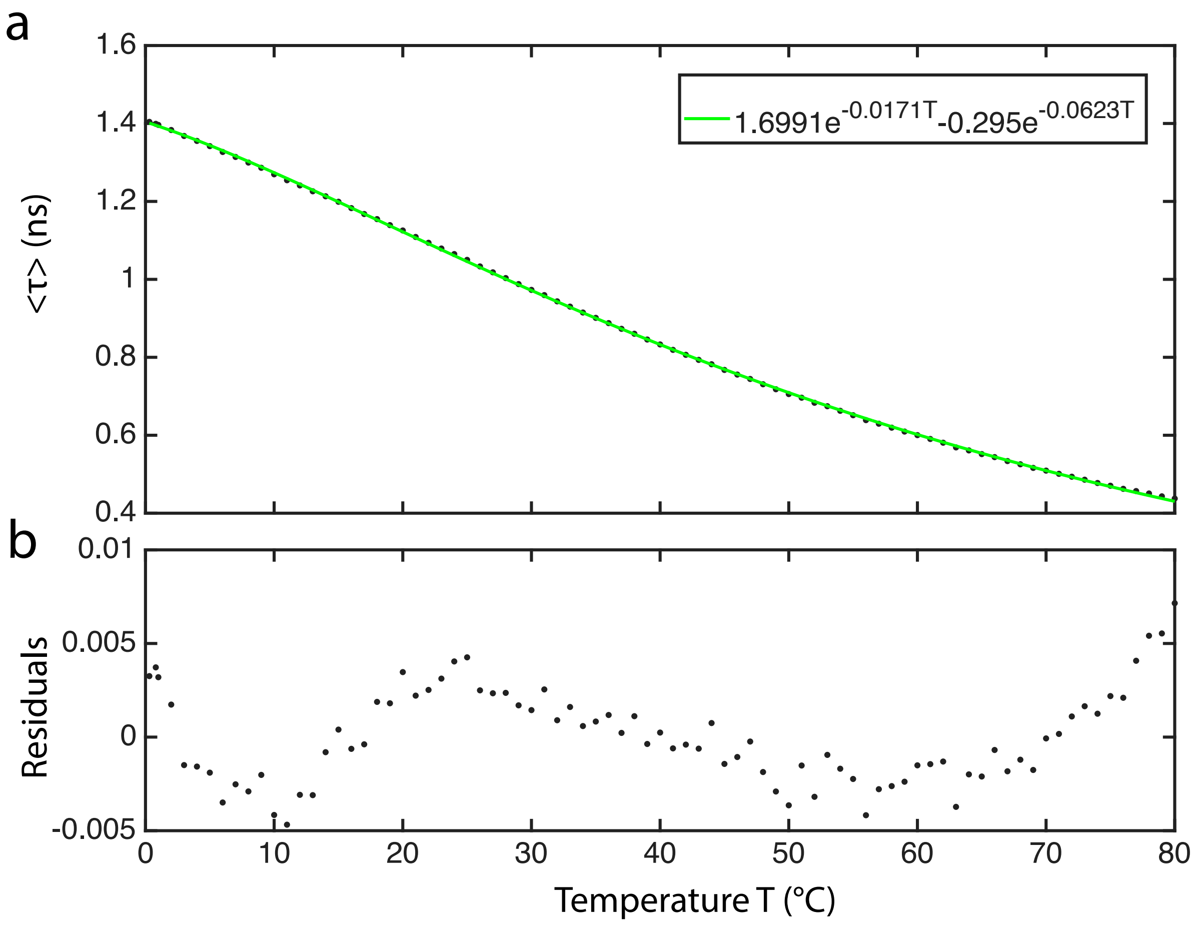


**Supplementary Figure 7: Biexponential regression to fluorescence lifetime of Cy5 vs temperature**
a) Plot of mean excited state lifetime ($\tau$) of Cy5 in cell culture media (black dots) measured at an Abberior Expert line confocal laser scanning microscope against temperature (N=4) together with a biexponential fit of the data (green line) b) Residuals from the fit (black dots).

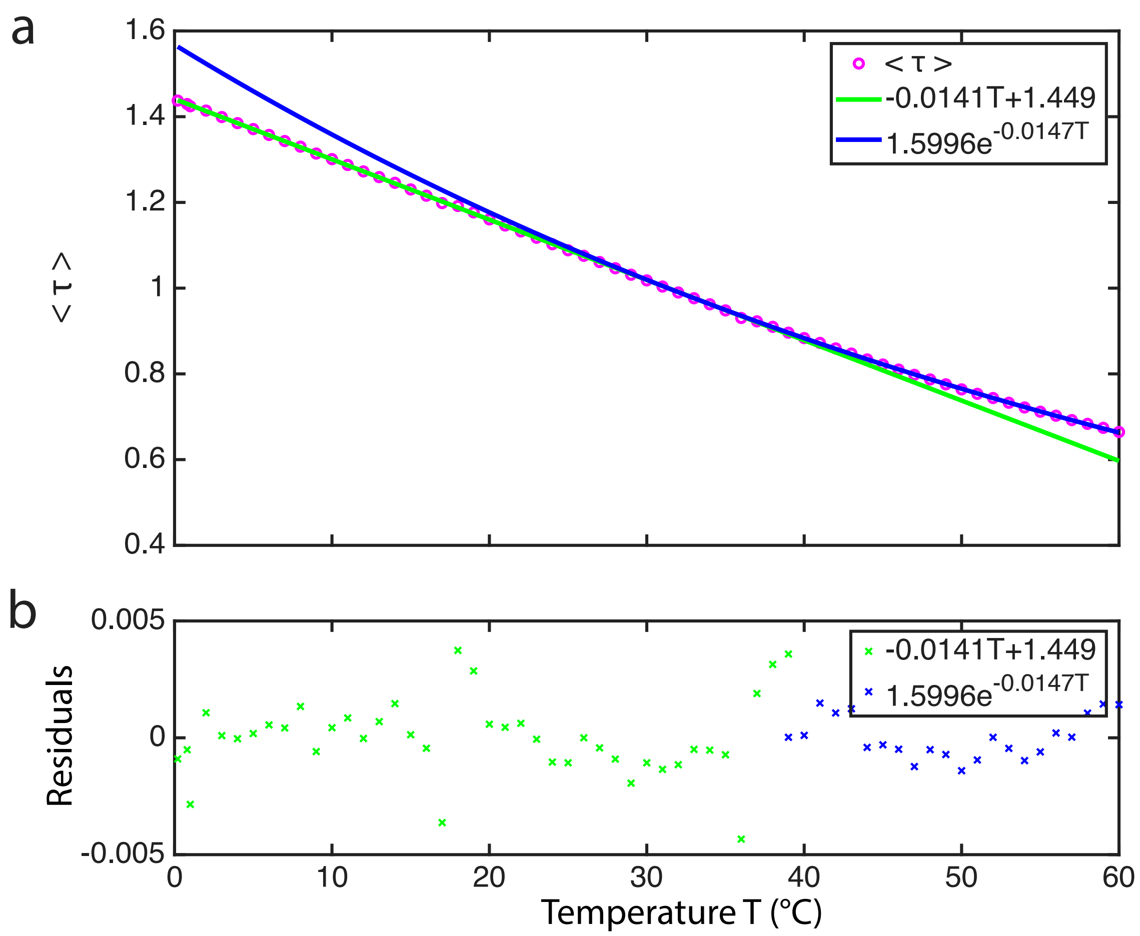


**Supplementary Figure 8: Excited state lifetime of Cy5 in cell culture buffer as function of temperature**

a) Excited state lifetime of Cy5 in a cell culture buffer (DMEM with 10% FCS) was measured over the temperature range 0-60°C on an Abberior Expert line CLSM. Shown is the mean excited-state lifetime ($\tau$) of Cy5 (circles) over temperature from N=3 independent experiments together with a linear function fitted to the data range from 0-39°C (green line) and a monoexponential function fitted to the temperature range 40°C-60°C (blue line). b) Residuals in both fit-regimes: linear (green crosses), monoexponential (blue crosses).

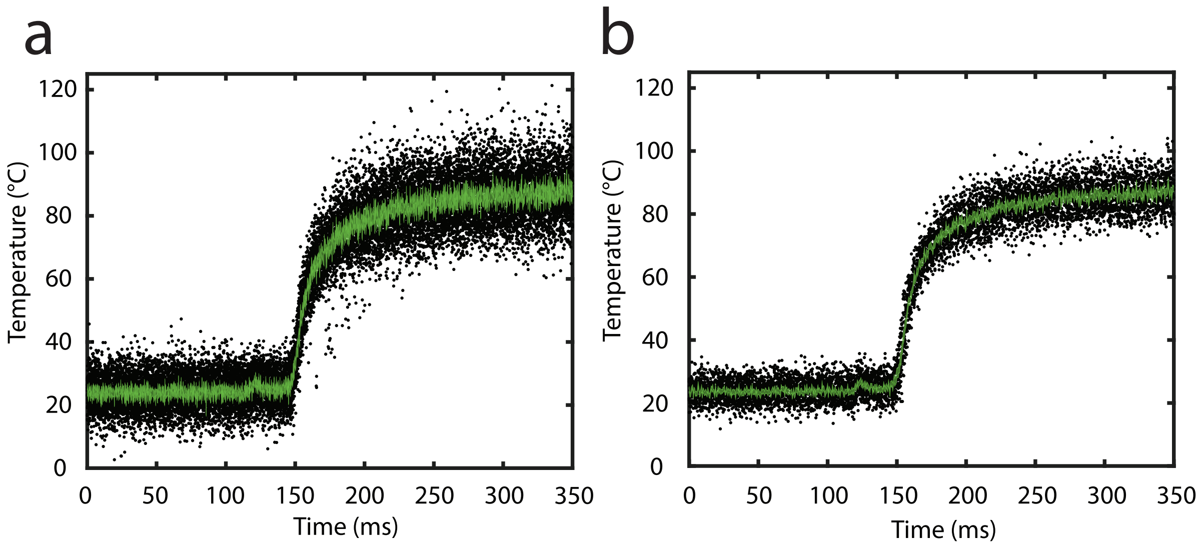


**Supplementary Figure 9: Temporal response of infrared laser heating sampled at 200 or 500 µs**

Temperature course upon turning-on 1470-nm laser illumination (110 mW), measured in the center of a flat-top laser beam of Ø = 160 µm. Black dots: Accumulated data points from n=21 experiments; green lines: mean ± s.e.m. a) sampling rate: 200 µs; b) 500 µs

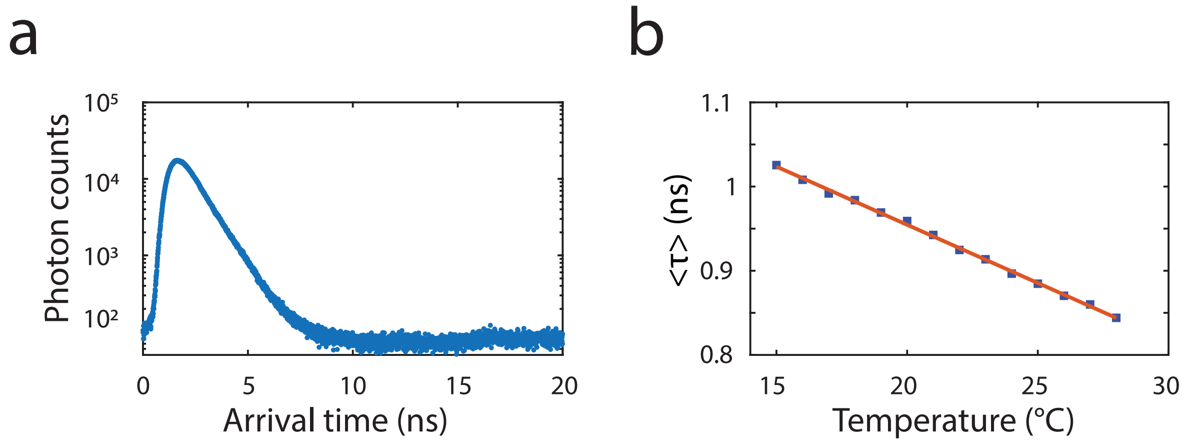


**Supplementary Figure 10: Fluorescence of Cy5 induced by a pulsed 775-nm laser**

a) Lifetime histogram of 6 µM Cy5 in water excited by a doughnut-shaped 775-nm laser at 23 mW b) Plot of mean excited state lifetime ($\tau$) of 6 µM Cy5 in water (black symbols) against temperature (N=2) together with a linear regression of the data (red line)
